# Amyloid precursor protein dosage normalization rescues neurogenesis and Alzheimer’s Disease phenotypes associated with Down Syndrome

**DOI:** 10.1101/2025.07.30.667706

**Authors:** Deepika Patel, Karen Rakowiecki, Orly Lazarov

## Abstract

Down Syndrome (DS) is the most abundant genetic form of mental retardation. It is caused by the triplication of partial or complete human chromosome 21 (HSA21). The molecular mechanisms causing it are not fully understood. Previous studies identified “Down syndrome Critical Region” (DSCR) genes that are essential or sufficient for the development of DS. However, these studies are largely inconclusive, due, in part, to the reliance on a small number of epidemiological cases. Amyloid precursor protein (*APP*) resides on HSA21 and is triplicated in DS. APP plays a role in developmental and post-natal neurogenesis, but is not thought to be part of the DSCR. The role of APP overdose in cortical malformation and cognitive impairments in DS is unknown. Mutations in *APP* cause familial Alzheimer’s disease (FAD). However, whether APP overdose is sufficient for the development of Alzheimer’s disease (AD) in DS is not fully understood. Here, we addressed the role of APP overdose in neuronal development and AD pathology. Using CRISPR/Cas9 gene editing, we eliminated one copy of *APP* from Down Syndrome-derived induced iPSCs DS APP(+/+/-) and examined the effect on neurogenesis, AD-related pathology and the expression levels of genes on HSA21 that are implicated in DS, neurodegeneration and inflammation.

## INTRODUCTION

Down Syndrome (DS) is a genetic disorder that is caused by the triplication of human chromosome 21 (HSA21)^1^. It is manifested by alterations in brain development, leading to brain dysfunction and intellectual disability^2,3^. However, how triplication of HSA21 induces these alterations is not fully understood. Previous attempts to narrow down genes that are essential or sufficient for the development of DS yielded several versions of the “Down syndrome Critical Region” (DSCR)^4^. These studies are largely inconclusive, due, in part, to the reliance on a small number of epidemiological cases. Amyloid precursor protein (*APP*) resides on HSA21, is triplicated in DS, but was not thought to be part of the DSCR. Notably, previous reports suggested several roles of APP in neurogenesis^5^, including the regulation of neural progenitor cell proliferation in the post-natal brain in mouse^6,7^ and human progenitors^8^. Mutations in *APP* cause familial Alzheimer’s disease (FAD)^9^. A large proportion of DS patients develop Alzheimer’s disease (AD)^10^. However, whether *APP* overdose is sufficient for the induction of brain malformation or the development of Alzheimer’s disease (AD) in DS is not fully understood. Previous studies suggested that *APP* overdose underlies increased b-amyloid (Ab) production, altered Ab_42_/_40_ and deposition of the pyroglutamate (E3)-containing amyloid aggregates, but not for tau-linked AD phenotypes^11^. Here, we addressed the role of APP overdose in neuronal development in DS. Using CRISPR/Cas9 gene editing, we eliminated one copy of *APP* from Down Syndrome-derived induced iPSC DS APP(+/+/-) and examined the effect on neurogenesis and AD-related pathology. We observed that the correction of APP dose rescues impairments in proliferation of neural progenitor cells and in differentiation of neural precursors and neurons. In addition to amyloid processing, we show that APP overdose increases tau expression and correcting APP rescues this increase. Notably, correction of APP does rescued overdose of other genes on HSA21, suggesting that it acts upstream of these genes. Together, this study suggests that APP overdose drives both brain maldevelopment and AD pathology, and that part of these effects can be attributed to gene interactions and linkage.

## RESULTS

### Down Syndrome iPSC - derived neural cells exhibit reduced proliferation and premature neuronal differentiation

We first examined the course of neuronal differentiation in DS using DS- and healthy control- (Ctrl) derived iPSC lines (**Figure 1A**). iPSC lines were differentiated into neural progenitor cells (NPCs) followed by neural precursor cells and forebrain neurons using neural induction method (**Figure 1B**). We assessed cell proliferation of NPCs using Click-It EdU assay. We observed a decrease in the proliferative capacity of DS NPCs compared to Ctrl, as indicated by a significantly reduced percentage of EdU+ cells in DS NPCs (**Figure 1C-D**). Further, the trajectory of neuronal differentiation progression was assessed by quantifying the expression of SOX2, DCX, and bIII-Tubulin in the cultures. As expected, immunofluorescence studies assessing SOX2 expression in NPCs, precursor and forebrain neurons revealed a decrease in SOX2+ cells across differentiation stages. However, DS iPSCs showed substantial increase in SOX2+ cells compared to Ctrl at each differentiation stage. (**Figure 1E-F**). In Ctrl lines, DCX expression peaked at the precursor stage. However, compared to Ctrl, a significant increase in the number of DCX+ cells was observed at each stage of neurogenesis (**Figure 1G-H**). Lastly, a significant increase in the number of b-III-tubulin+ precursors and neurons was observed in the DS lines compared to Ctrl (**Figure 1I-J**). To further validate the neuronal differentiation phenotypes, we examined the expression levels of these proxies by Western blot analysis (**Figure 2A**). Results revealed significant alterations in the expression of neuronal proxies during the differentiation of DS iPSCs-derived neural cells compared to Ctrl. Specifically, and in agreement with the immunofluorescence analysis, we observed increased expression of Sox2 in NPCs and precursors of DS lines compared to Ctrl (**Figure 2A,E**), increased DCX (**Figure 2A,D**) and β-III-tubulin (**Figure 2A,C**) expression in NPCs, precursors and 10 weeks neurons in DS lines (**Figure 2A-E**), and increased neurofilament heavy chain in 10 week DS neurons (**Figure 2A,B**), indicative of enhanced neuronal maturation. Collectively, these results indicate that DS iPSCs-derived neural cells display defects in proliferation and enhanced premature neuronal differentiation, mirroring neurodevelopmental abnormalities observed in Down Syndrome^12,13^.

**Figure 1.**
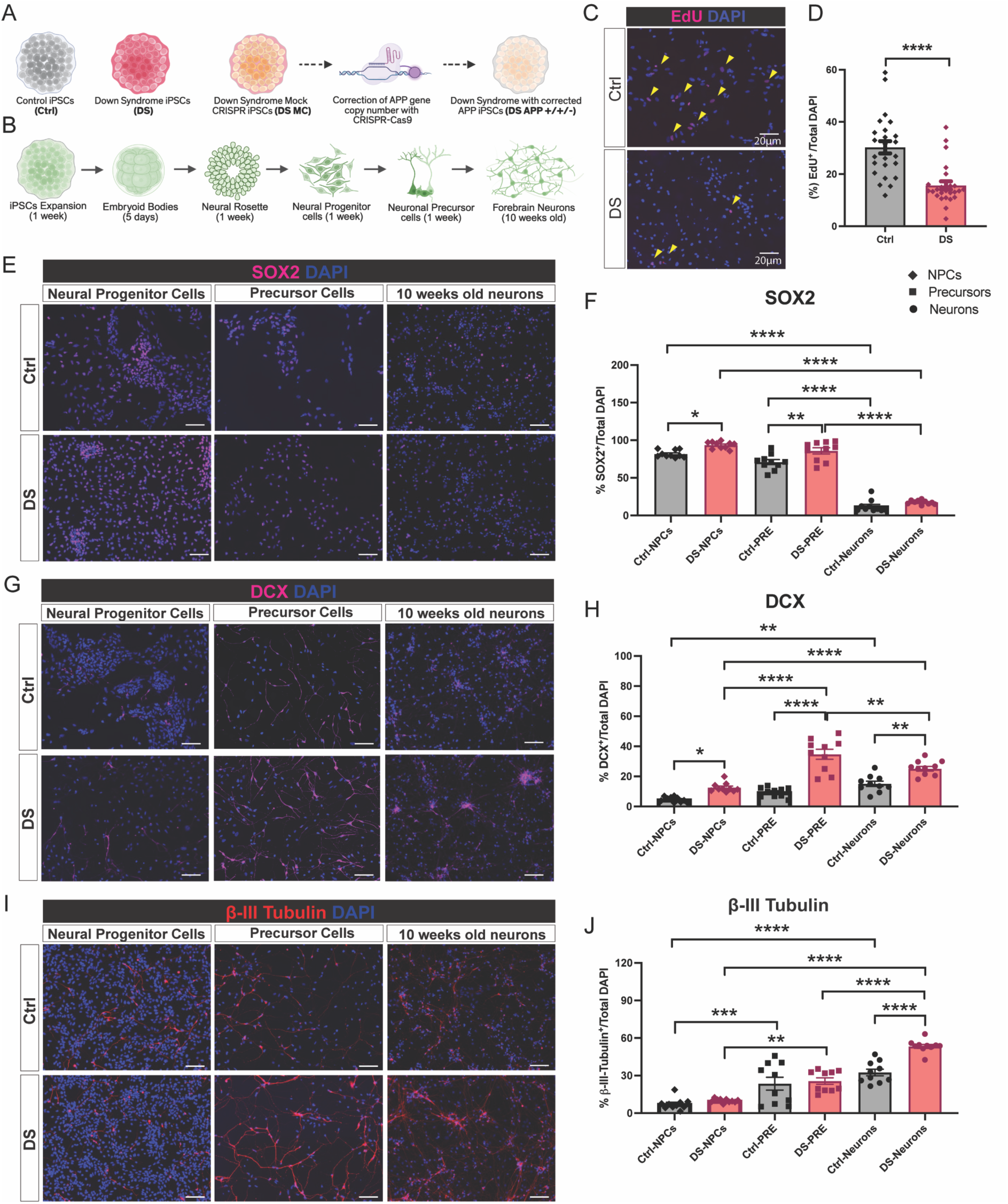
Developmental neurogenesis in control and Down Syndrome iPSCs-derived neural cells. (A) Schematic of patient-derived pluripotent stem cells (iPSCs) experimental groups. Control iPSCs (Ctrl), Down syndrome iPSCs (DS), gene-corrected DS iPSCs (DS APP*+/+/-*) and a mock CRISPR control (DS MC). (B) Timeline of differentiation of iPSCs to forebrain neurons, indicating the key stages: iPSCs expansion, embryoid body formation, neural rosette induction, neural progenitor cell (NPCs), precursor cells (PRE), and 10-week-old forebrain neurons. (C) Representative EdU immunocytochemistry images showing proliferating (EdU+) neural progenitor cells derived from control and DS iPSCs, and co-stained with DAPI. Yellow arrowheads mark EdU+ cells. Scale bars: 20µm. (D) Quantification of EdU incorporation (%EdU/total DAPI) shows significantly decreased proliferation in DS neural progenitor cells compared to controls. (E, G, I) Representative immunofluorescence images of SOX2 (magenta), DCX (magenta), b-III-tubulin (red) and DAPI (blue) in neural progenitor cells, precursor cells, and 10-week-old neurons from control and DS groups. Scale bars: 20µm. (F, H, J) Quantification of SOX2+, DCX+, and β-III-tubulin+ cells as a percentage of total DAPI+ nuclei across differentiation stages, demonstrates enhanced neuronal differentiation in DS lines. Data represented as mean ± SEM with individual points shown. Data analyzed by unpaired t-test (D), two-way ANOVA with Sidak’s multiple comparisons test (F, H, J), *p < 0.05, **p < 0.01, ***p < 0.001, and ****p < 0.0001.

**Figure 2.**
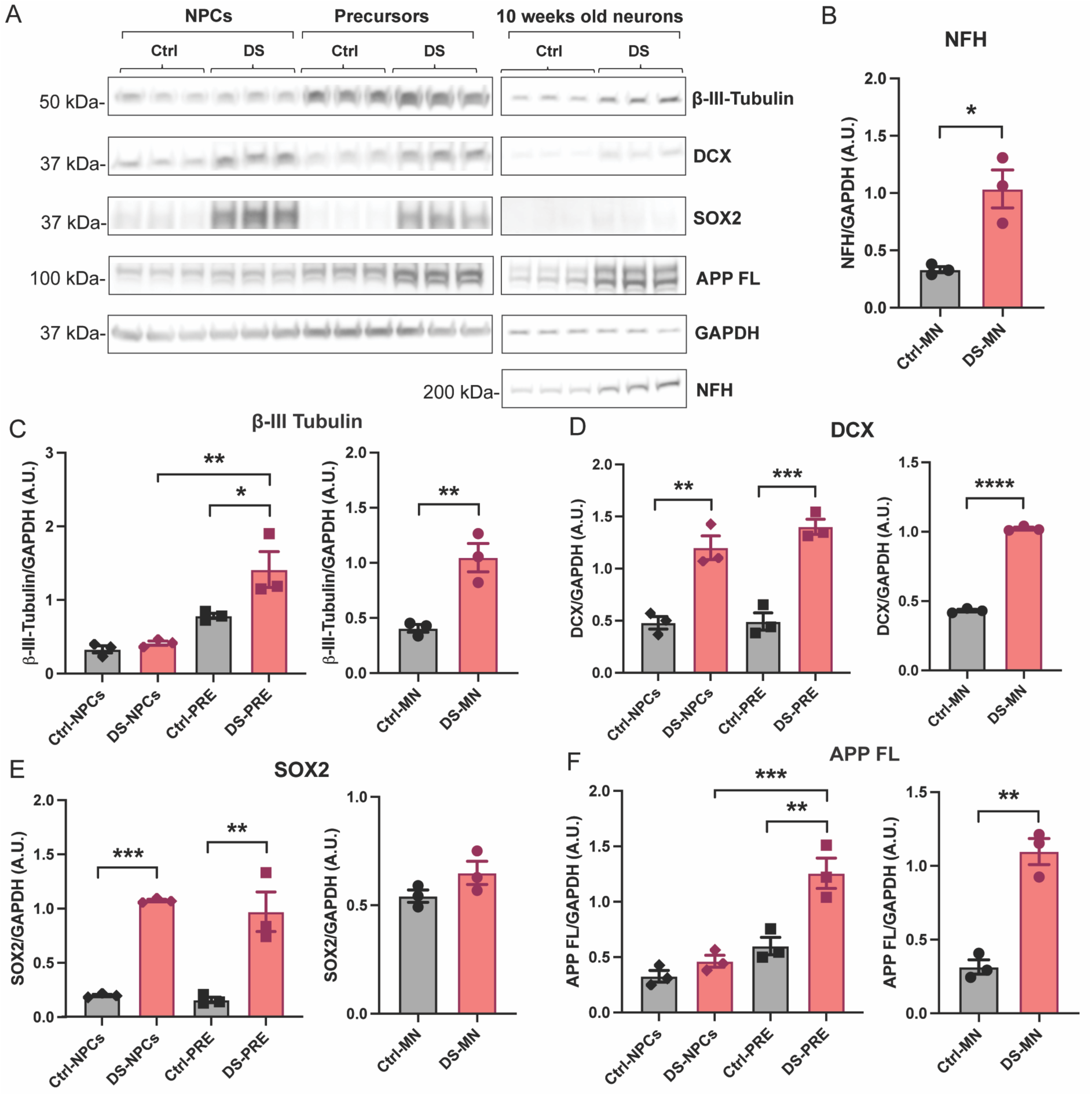
Altered protein expression profile of key neurogenesis signals in DS iPSCs-derived neural populations. (A) Expression levels of β-III-Tubulin, DCX, SOX2, full-length APP (FL-APP) and neurofilament heavy chain (NFH) in NPCs, PRE, and 10-week-old neurons (MN) derived from Control (Ctrl) and Down Syndrome (DS) iPSCs, as detected by Western blot analysis. (B) Quantification of NFH normalized to GAPDH in 10-week-old neurons indicates a significant increase in DS-derived neurons compared to control. (C, D, E, F) Quantification of β-III-Tubulin, DCX, SOX2, APP full-length expression normalized to GAPDH in NPCs, precursor cells (left) and neurons differentiated from control and DS groups (right) supports increased neuronal differentiation in the DS group. Data represented as mean ± SEM with individual data points shown. Data analyzed by unpaired t test (neurons), one-way ANOVA with Tukey’s multiple comparisons test (NPCs and precursor cells data), *p < 0.05, **p < 0.01, ***p < 0.001, and ****p < 0.0001.

### Enhanced amyloidogenic APP processing and tau accumulation in Down Syndrome iPSCs-derived neurons

Next, we examined the development of AD -related neuropathology in our DS and Ctrl lines. We first assessed the protein levels of FL-APP (full length) at differentiation stages of cortical neurogenesis. FL-APP was upregulated in DS precursors and neurons compared to Ctrl counterparts (**Figure 2A, F)**, aligning with the gene dosage effect of HSA21. In addition, levels of BACE1, APP-carboxyl terminal fragments (APP-CTFs, **Figure 3C**), total tau (Tau-5, **Figure 3E**) and phosphorylated tau (AT8, **Figure 3D**) were upregulated in protein lysate of 10 weeks old DS iPSCs-derived neurons compared to Ctrl (**Figure 3A-F**). The ratio of phosphorylated to total tau (AT8/Tau5) did not differ significantly (**Figure 3F**). Correspondingly, ELISA measurements revealed a marked increase in secretion of both Aβ_40_ (**Figure 3G**) and Aβ_42_ (**Figure 3H**) peptides in DS neurons, albeit the Ab_42_/Ab_40_ ratio remained unchanged between groups (**Figure 3I**). Together, these results suggest enhanced amyloid and tau pathology in the DS neurons, and illustrate that DS iPSCs-derived neurons recapitulate key molecular hallmarks of AD, including increased amyloidogenic processing and tau accumulation, providing a robust *in vitro* model for investigating AD pathology in the context of DS.

**Figure 3.**
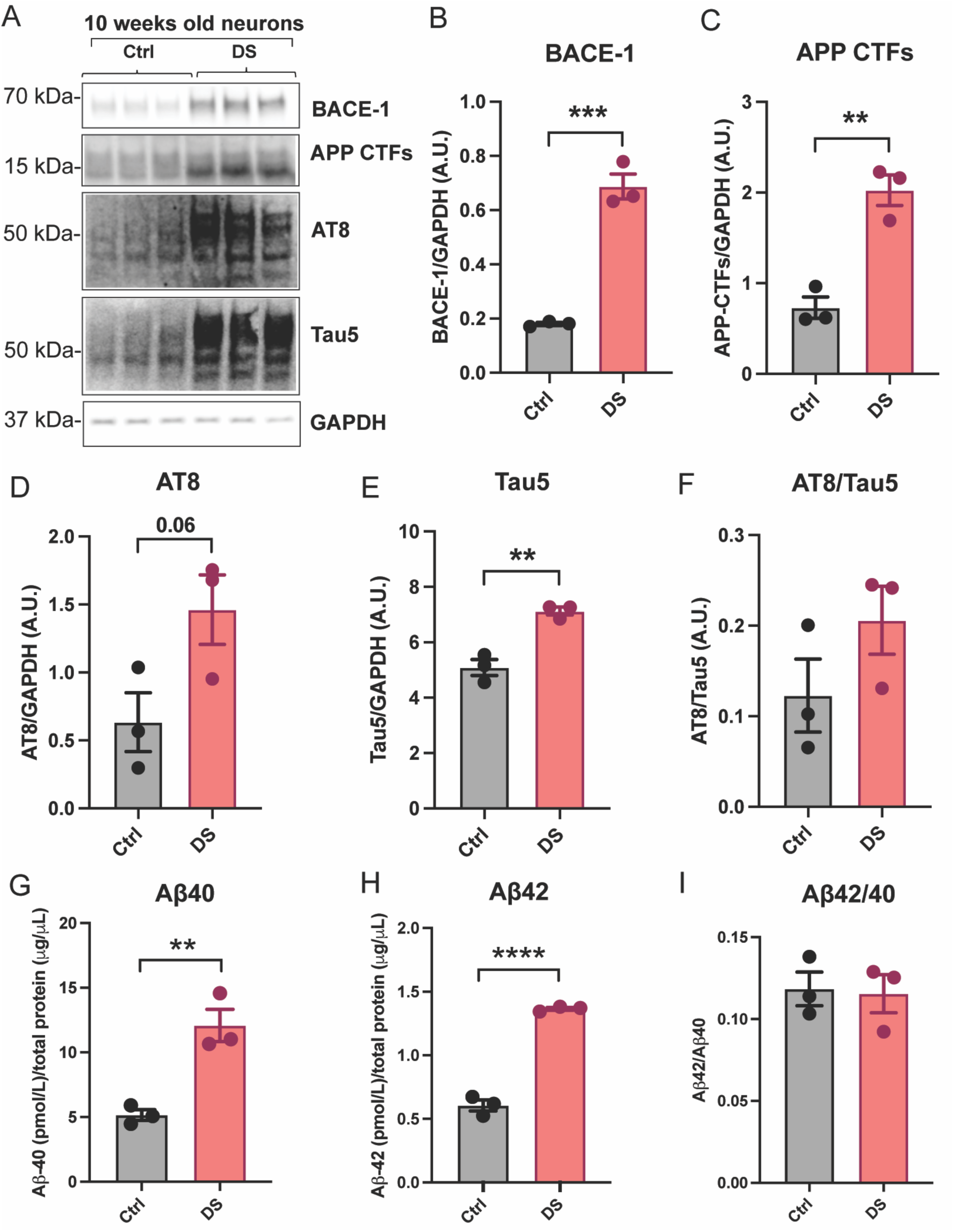
Increased amyloidogenic processing of APP and tau in Down Syndrome iPSCs-derived neurons, recapitulating selected Alzheimer’s disease-related phenotypes. (A) Increased expression of BACE-1, APP C-terminal fragments (CTFs), phosphorylated tau (AT8), and total tau (Tau5) in protein lysate of 10-week-old neurons derived from DS iPSCs compared to Ctrl, as detected by Western blot analysis. (B-E) Semi-quantification of BACE-1, APP CTFs, AT8, Tau5 levels normalized to GAPDH in Ctrl and DS groups. (F) AT8/Tau5 ratio indicates no significant difference between DS and Ctrl groups. (G-H) Quantification of secreted Ab_40_ and Ab_42_ in DS neuronal cultures compared to controls using ELISA. (I) The Ab_42_/Ab_40_ ratio is comparable between genotypes. Data represented as mean ± SEM with individual data points shown. Data analysed by unpaired t test, **p < 0.01, ***p < 0.001, and ****p < 0.0001.

### Coordinated upregulation of chromosome 21 encoded and stress-associated proteins in Down Syndrome neurons

Next, we asked whether alterations in expression of additional genes on HSA21 would be apparent in our iPSC-derived DS neurons. Therefore, we examined the expression of selected genes that are implicated in inflammation, neurodegeneration and neurogenesis. Specifically, DYRK1A, BACE2, SOD1 and S100b. Quantitative analysis of protein expression in 10-week-old iPSCs-derived neurons revealed the upregulation of DYRK1A (**Figure 4A-B),** BACE-2 (**Figure 4A, C**) and SOD1 **(Figure 4A, D)** in the DS neurons compared to Ctrl. Notably, DS neurons showed a robust increase in S100β dimer levels (**Figure 4A, E**), while monomer levels remained unchanged between groups (**Figure 4A, F**). Together, these findings highlight the pervasive impact of chromosome 21 trisomy on gene and protein expression profiles, implicating these molecular changes in disrupted cellular homeostasis and heightened stress responses in DS-derived neural populations.

**Figure 4.**
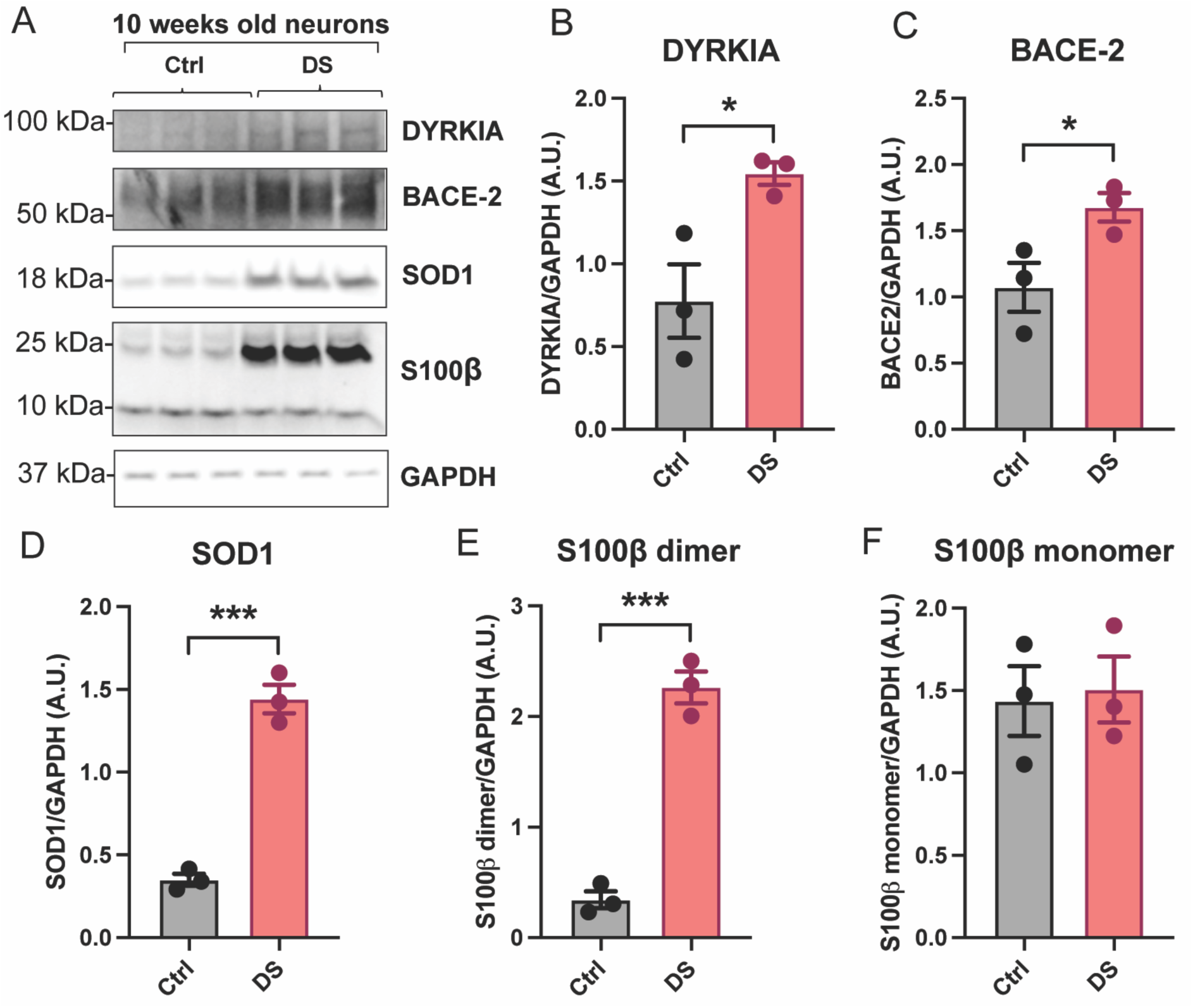
Upregulation of chromosome 21 genes and stress response proteins in Down Syndrome iPSCs-derived neurons. (A) Protein expression levels of DYRK1A, BACE-2, SOD1, and S100b in 10-week-old neurons differentiated from Ctrl and DS iPSCs, as detected by Western blot analysis. (B-F) Quantification of DYRK1A, BACE-2, SOD1, and S100b expression, normalized to GAPDH. Data represented as mean ± SEM with individual data points shown. Data analysed by unpaired t-test, *p < 0.05, and ***p < 0.001.

### Normalization of *APP* copy number rescues neurogenesis deficits in Down Syndrome iPSCs-derived neurons

Having established the phenotypes of DS iPSC-derived neurogenesis and neurons, we examined the hypothesis that *APP* gene copy number and resultant overdose contributes to deficits in neurogenesis and the development of AD pathology in DS. For that, we utilized CRISPR/Cas9 technology to eliminate one copy of *APP* from DS-derived iPSCs (DS APP*+/+/-*) and to generate a mock CRISPR-Cas9 DS control (DS MC) (**Figure 5A**). Driven by this hypothesis, we investigated whether the restoration of normal *APP* gene dosage in DS iPSCs rescued neurogenesis deficits. Our results showed that correction of *APP* gene dosage in DS iPSCs-derived neural cultures leads to rescue of neurogenesis deficits. Specifically, DS APP*+/+/-* cells exhibited a marked increase in the percentage of proliferating NPCs, as shown by greater EdU incorporation compared to DS MC (**Figure 5B-C**). Immunofluorescence quantification further demonstrated restoration of SOX2+ cell number at all three stages of differentiation (**Figure 5D-E**) and normalization of the number of DCX+ cells at neuronal precursor and neuronal stages in DS APP*+/+/-* group (**Figure 5F-G**). Moreover, the number of β-III-tubulin+ neurons was normalized in DS APP*+/+/-* compared to DS MC (**Figure 5H-I**).

**Figure 5.**
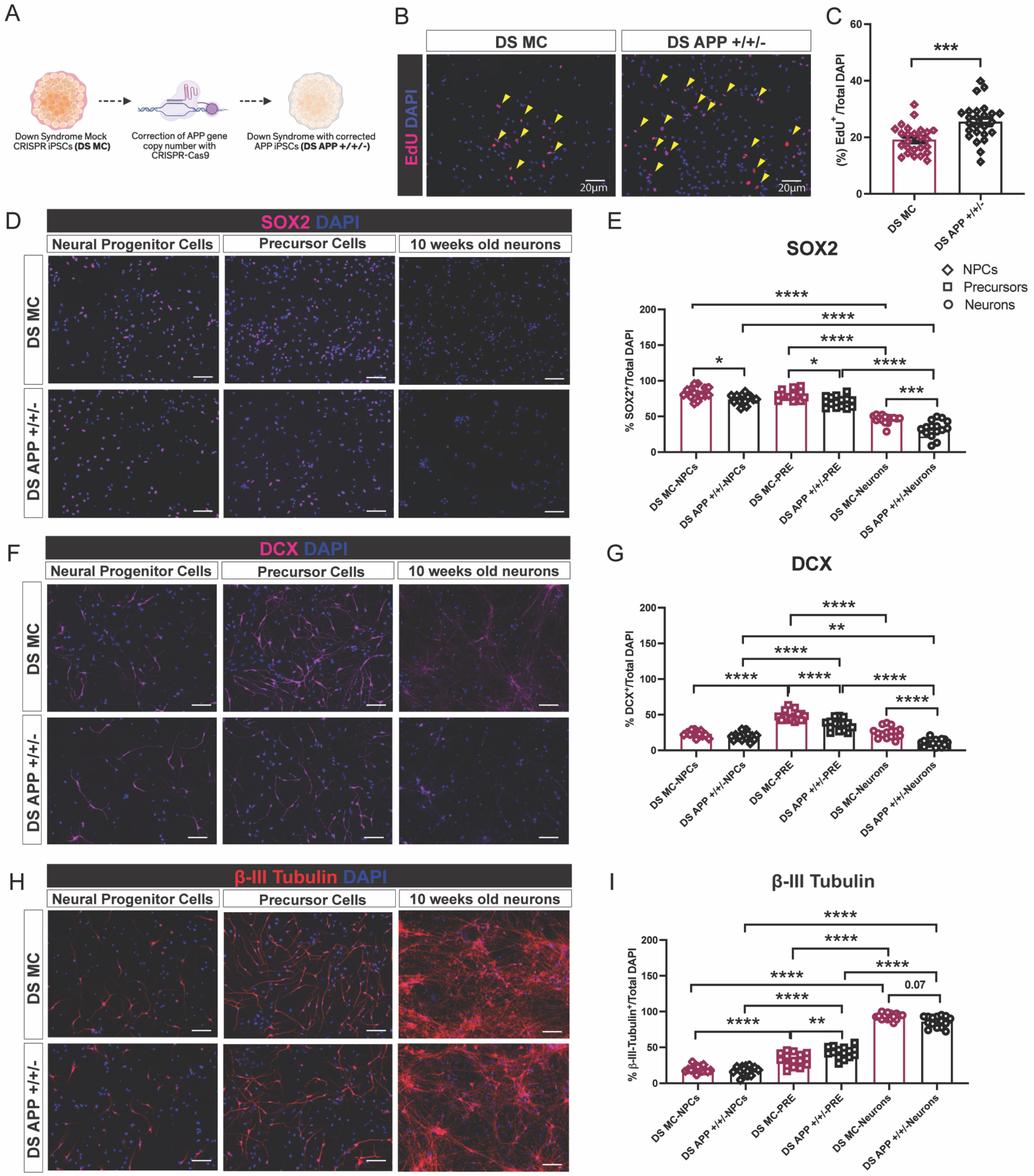
Correction of *APP* gene dosage rescues proliferation and neuronal differentiation defects in Down Syndrome iPSCs-derived cells. (A) Schematic depicting the manipulation of DS iPSCs with either mock CRISPR editing (DS MC) or APP gene copy correction (DS APP*+/+/-*). (B) Representative immunofluorescence images showing EdU incorporation (red) and DAPI (blue) in NPCs from DS MC and DS APP+/+/- lines. Yellow arrowheads indicate EdU+ proliferating cells. Scale bar: 20µm. (C) Quantification of EdU+ cells as a percentage of total DAPI-labelled nuclei, suggesting significantly increased proliferation in DS APP*+/+/-* NPCs compared to DS MC. (D, F, H) Representative images of SOX2 (magenta), DCX (magenta), β-III-Tubulin (red), and DAPI (blue) immunostaining in NPCs, precursor cells, and 10 weeks old neurons derived from DS MC and DS APP*+/+/-* iPSCs. Scale bars: 20µm. (E, G, I) Quantification of SOX2+, DCX+, and β-III-Tubulin+ cells (% of total DAPI) across differentiation stages. Data represented as mean ± SEM with individual points shown. Data analyzed by unpaired t-test (C), two-way ANOVA with Sidak’s multiple comparisons test (E, G, I), *p < 0.05, **p < 0.01, ***p < 0.001, and ****p < 0.0001.

To further validate these findings, we examined the expression levels of these proxies in protein lysates of these lines. Normalization of *APP* gene dosage in DS iPSCs-derived neural cultures validated the robust rescue of aberrant neurogenesis proxies (**Figure 6A**). Western blot analyses demonstrated no significant change in NFH protein levels in both the groups (**Figure 6A-B**). DS APP*+/+/-* NPCs, precursors and neurons exhibited markedly reduced levels of β-III-tubulin (**Figure 6A, C)**, DCX (**Figure 6A, D**), and SOX2 (**Figure 6A, E)** compared to DS MC controls. Collectively, these results indicate that restoring *APP* gene dosage corrects proliferative and differentiation abnormalities in DS iPSCs-derived neural cells.

**Figure 6.**
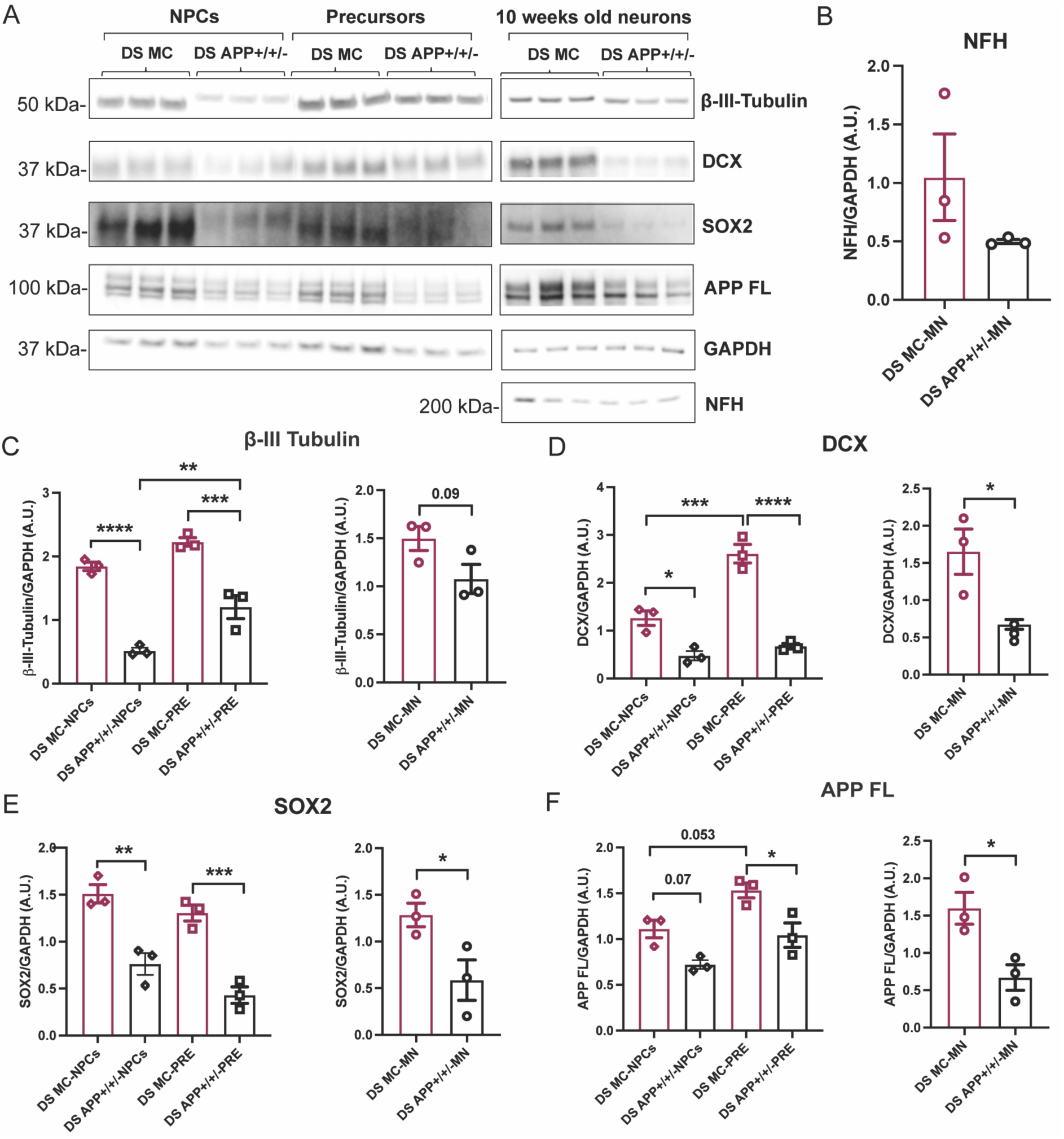
*APP* dosage correction reverses aberrant neurogenesis proxy expression in Down Syndrome iPSCs-derived cells. (A) Expression levels of b-III-Tubulin, DCX, SOX2, APP full-length (APP FL) and neurofilament heavy chain (NFH) across NPCs, precursor cells, and 10 week-old-neurons derived from DS iPSCs with DS MC and DS APP+/+/- groups, as detected by Western blot. (B) Quantification of NFH expression normalized to GAPDH in 10-week-old neurons. (C, D, E, F) Quantification of b-III-Tubulin, DCX (doublecortin), SOX2, APP full-length expression normalized to GAPDH in NPCs and precursor stages (left) and in neurons differentiated from APP protein levels in DS APP*+/+/-* and mock control iPSC lines. Data represented as mean ± SEM with individual data points shown. Data analyzed by unpaired t-test (neurons), one-way ANOVA with Tukey’s multiple comparisons test (NPCs and precursor cells data), *p < 0.05, **p < 0.01, ***p < 0.001, and ****p < 0.0001.

### Rescue of Alzheimer’s disease molecular phenotypes by APP dosage normalization in Down Syndrome neurons

Next, we asked whether the normalization of *APP* copy number and dose, rescues the increase in APP processing and subsequent AD-linked pathology. We observed that FL-APP protein levels were reduced at all stages in DS APP*+/+/-* lines compared to DS Mock (**Figure 6A, F)**, confirming successful genetic correction. Furthermore, normalization of *APP* gene dosage in DS iPSCs-derived neurons profoundly attenuates alterations in AD-related signals. Specifically, we observed significantly reduced levels of BACE-1 in DS APP*+/+/-* neurons compared to DS MC (**Figure 7A-B**). There was no change in levels of APP-CTFs between the genotypes (**Figure 7A, C)**. Both AT8 (**Figure 7A, D**) and Tau5 **(Figure 7A, E)** were markedly diminished in DS APP*+/+/-* neurons compared to DS MC, demonstrating normalization of tau metabolism. The ratio of AT8/Tau5 remained unchanged (**Figure 7F**), suggesting that *APP* correction primarily reduces overall tau levels rather than alters phosphorylation stoichiometry. Levels of secreted Aβ_40_ (**Figure 7G**) and Aβ_42_ (**Figure 7H**) were robustly decreased in DS APP*+/+/-* neurons. Ab_42_/Ab_40_ ratio showed trend, albeit not statistically significant (**Figure 7I**). Collectively, these results demonstrate that restoring *APP* dosage levels in DS iPSCs-derived neurons rescues key AD-associated biochemical phenotypes.

**Figure 7.**
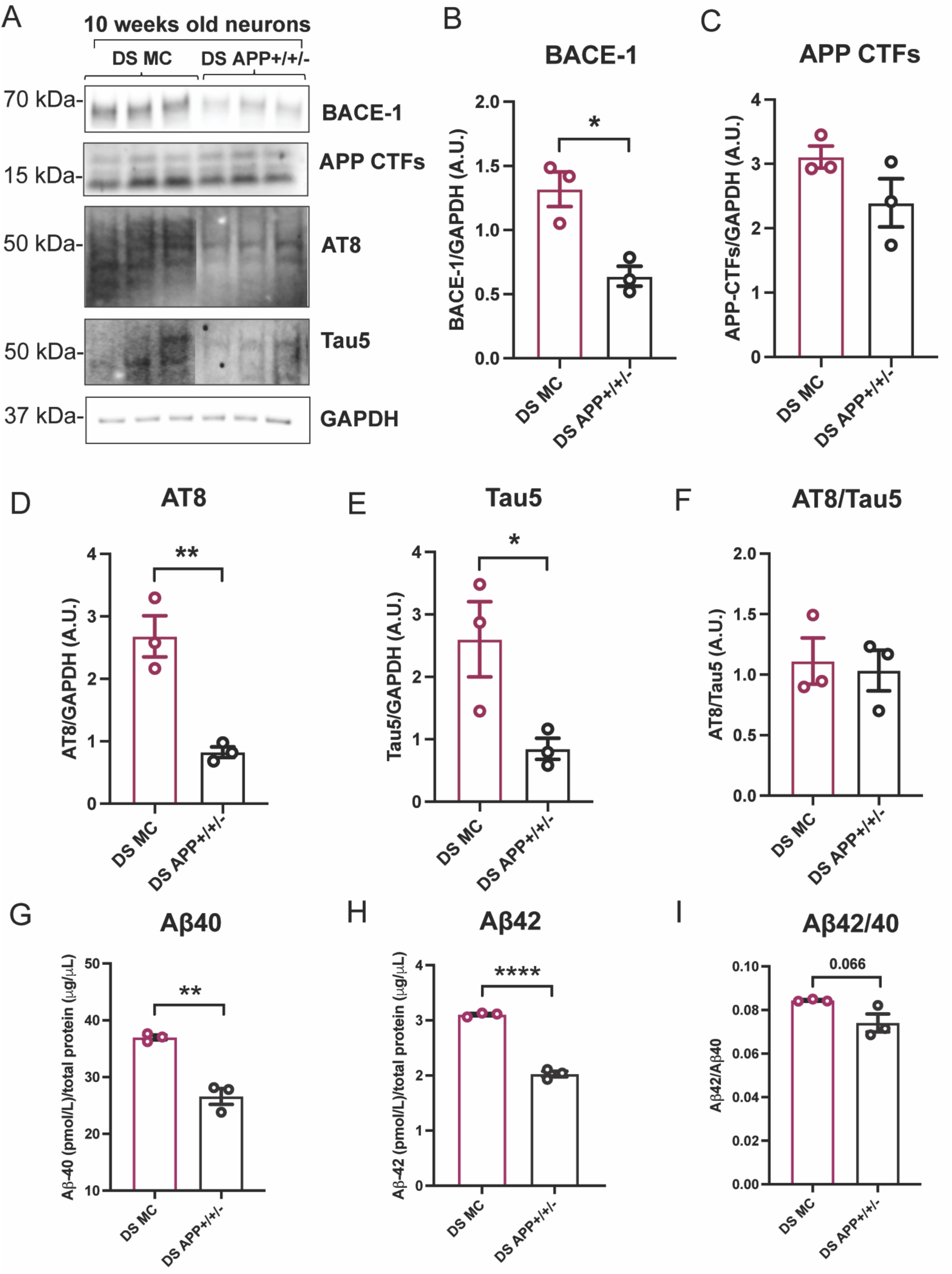
APP Dosage Correction attenuates Alzheimer’s disease related pathology in Down Syndrome iPSCs-derived neurons. (A) Expression levels of BACE-1, APP C-terminal fragments (APP CTFs), phosphorylated tau (AT8), total tau (Tau5) in protein lysate of 10 week old neurons derived from DS MC and DS APP+/+/- iPSC lines, as examined by Western blot. (B-E) Quantification of BACE-1, APP CTFs, AT8, Tau5 levels normalized to GAPDH in DS MC and DS APP*+/+/-* groups. (F) The ratio of phosphorylated- to total- tau (AT8/Tau5) is comparable between genotypes. (G, H) Quantification of secreted Ab_40_ and Ab_42_ in conditioned media of DS APP*^+/+/-^* vs DS MC neuronal cultures using ELISA. (I) Ab_42_/Ab_40_ ratio following *APP* dosage correction. Data represented as mean ± SEM with individual data points shown. Data analyzed by unpaired t-test, *p < 0.05, **p < 0.01, and ****p < 0.0001.

### APP gene normalization diminishes Chromosome 21 gene dosage effects and stress pathways in Down Syndrome neurons

Finally, we examined whether the normalization of *APP* gene dosage in DS iPSCs-derived neurons had an effect on HSA21 genes that were observed to be upregulated (**Figure 4**). Examination of protein lysate of DS APP*+/+/-* and DS MC neurons revealed reduced expression of DYRK1A and BACE2 in DS APP*+/+/-* neurons compared to DS MC, with BACE2 levels being significant and DYRK1A trending but not significant (**Figure 8A, C**). Levels of SOD1 and S100β dimers and monomers were comparable between the genotypes (**Figure 8A, D-F**). These data indicate that *APP* gene dosage in DS neurons affects the dysregulation of genes on HSA21, underscoring the central regulatory role of APP in DS neuropathology.

**Figure 8.**
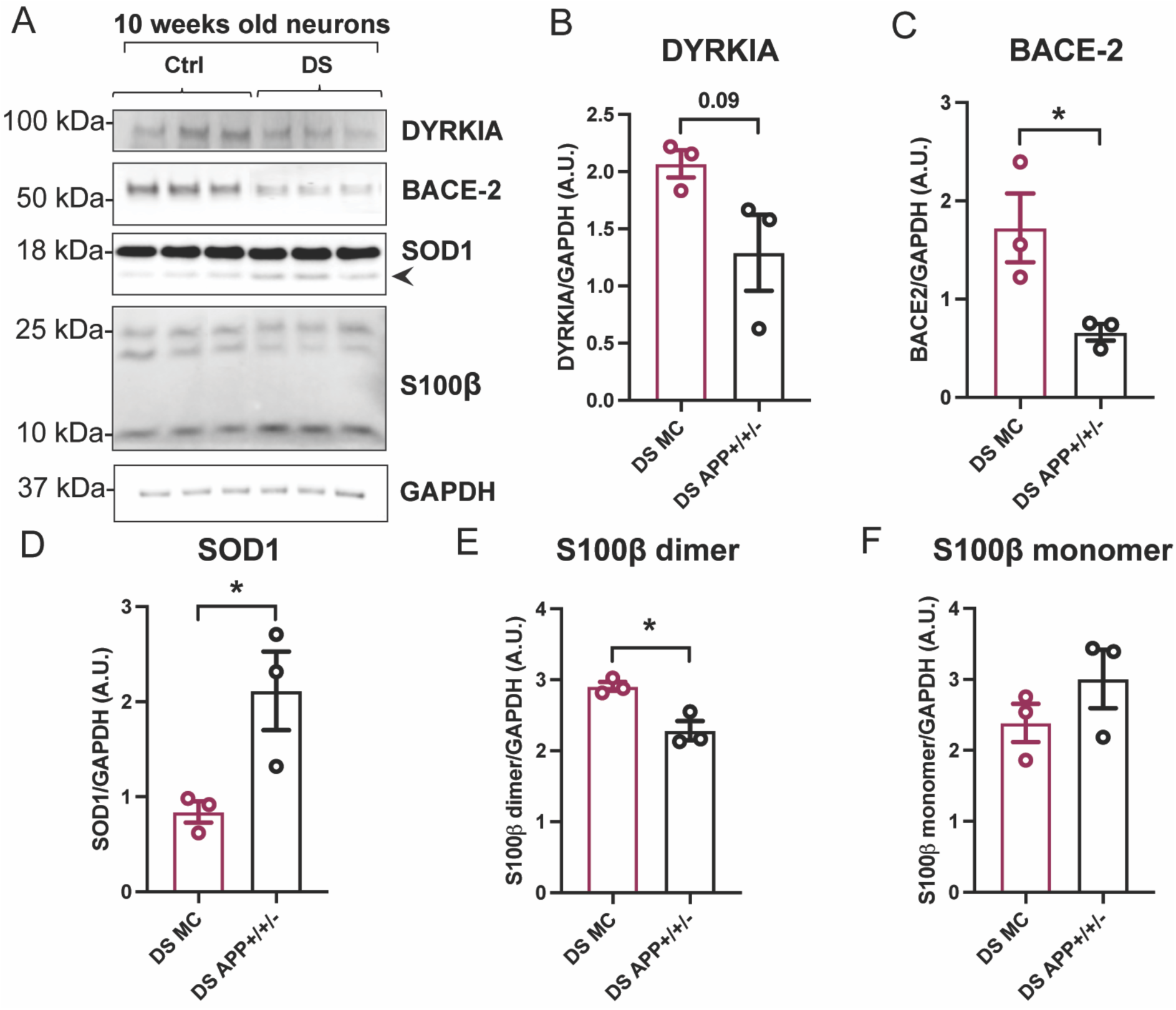
Correction of *APP* dosage rescues chromosome 21 gene and stress response protein dysregulation in Down Syndrome iPSCs-derived neurons. (A) Protein levels of DYRK1A, BACE-2, SOD1, and S100b in lysate of 10 weeks old neurons derived from DS MC and DS APP*+/+/-*, as detected by Western blot analysis. (B-F) Quantification of DYRK1A, BACE-2, SOD1, and S100b expression levels, normalized to GAPDH. Data represented as mean ± SEM with individual data points shown. Data analyzed by unpaired t-test, *p < 0.05.

## DISCUSSION

This study reveals several novel aspects. First, we show that APP overdose impairs NPCs proliferation. Interestingly, we observed upregulation of Sox2 in NPCs and precursor cells and increased number of cells expressing Sox2. On the other hand, we observed reduced proliferation of NPCs. Together, this may suggest enhanced stemness due to APP overdose. In addition, we observed that APP overdose induced premature differentiation of DS forebrain neurons. This may suggest that APP regulates more than one checkpoint along the neurogenic path. Further studies are warranted to determine if a specific metabolite of APP underlies one or more of these impairments. In addition, future studies should examine the implications for neuronal function. In that regard, we observed upregulation of tau species in DS neurons. Given the importance of tau to neuronal development, its upregulation may compromise neuronal maturation. Notably, we did not observe increased tau phosphorylation relative to total tau, suggesting that other factors may play a role in tau hyperphosphorylation. One possibility is that a two-hit, such as aging and APP overdose, would be required for the pathological acceleration of tau hyperphosphorylation. While we observed enhanced expression of Bace1 and enhanced secretion of Ab_42_ and Ab_40_ in DS neurons that were all rescued by APP dose correction, in contrast to previous studies^11^, we did not observe altered Ab_42_/Ab_40_ ratio. The DS MC and DS DS APP*+/+/-* lines developed in this study, provide a platform for the examination of the effect of APP overdose on many aspects of developmental neurogenesis and AD pathology that are yet to be examined in this setting. Limitation of our study include the 2D differentiation which does not allow full assessment of corticogenesis, and the lack of mixed cellular interactions with glia and microglia. Finally, the potential difficulty to facilitate amyloid deposition and neurofibrillary tangles in these cultures. Future studies should use these iPSC lines for the development of cortical organoids. Finally, we show that *APP* dose correction rescues alterations in additional genes on HSA21, suggesting functional interactions between these genes, and that APP governs these interactions. In summary, this study suggests that APP overdose causes impairments in neurogenesis and facilitates amyloidosis in DS.

## MATERIALS AND METHODS

### Human iPSCs Culture, NPCs differentiation and neuron maturation

The iPSC lines used in this study were obtained from repositories and collaborating laboratories. Control line Ctrl #1 (UWWC1-DS2U) and Down syndrome line DS #1 (UWWC1-DS1) were obtained from WiCell. The DS mock-corrected (MC) and APP gene-corrected (APP+/+/-) lines were graciously supplied by the Young-Pearse laboratory. Schematic representation of the experimental groups are depicted in **Figure 1A**.

Frozen vials from master stocks were thawed at 37°C and plated onto Matrigel hESC-Qualified Matrix-coated plates (Corning, 354277) in mTeSR Plus medium (STEMCELL Technologies, 100-0276), supplemented with 10 μM Y-27632 (ROCK inhibitor) during the first 24 hours post-thaw to enhance survival. The medium was refreshed daily. Individual colonies were mechanically dissected and passaged weekly. Unless otherwise noted, all reagents were purchased from STEMCELL Technologies.

Neural differentiation was performed following STEMCELL Technologies’ protocols, with key stages and validation illustrated in **Figure 1B**. To generate CNS-type neural progenitor cells (NPCs) from iPSCs, we employed the STEMdiff Neural Induction Medium with SMAD pathway inhibitors (SMADi) using an embryoid body (EB) formation approach. Briefly, confluent iPSCs were dissociated with Gentle Cell Dissociation Reagent and seeded at 3 x 10^6 cells per well into AggreWell 800 plates (preconditioned with Anti-Adherence Rinsing Solution), resulting in approximately 10,000 cells per microwell. The medium was partially replaced daily over five days. EBs were subsequently collected and plated onto poly-L-ornithine (PLO) (Millipore sigma, A-004-C)/laminin (Fischer scientific, 23-017-015)-coated 6-well plates to induce neural rosette formation. After 7 days, neural rosettes were isolated using STEMdiff Neural Rosette Selection Reagent and replated on PLO/laminin-coated plates for further NPCs expansion. NPCs were either banked or used directly for differentiation and expanded up to 4 passages.

For differentiation into neural precursors, NPCs were cultured for 7 days in STEMdiff Neural Progenitor Medium, then dissociated with accutase and plated in STEMdiff Forebrain Neuron Differentiation Medium for an additional week. Neural precursors were matured using the STEMdiff Forebrain Neuron Maturation Kit. All culture steps were performed on tissue culture-treated plastic plates, with 24-well glass-bottom plates used for imaging and immunocytochemical analyses. This differentiation protocol reliably produces validated NPCs, precursor cells, and 10 weeks old forebrain neurons suitable for downstream analyses.

### Immunocytochemistry

For immunocytochemistry (ICC) on cultured cells, culture media was removed, and cells were washed twice with 1X PBS. Cells were then fixed in 4% paraformaldehyde (PFA) prepared in 1X PBS for 15 minutes at room temperature (RT). Following fixation, cells were washed twice with 1X PBS and blocked in blocking solution consisting of 1X PBS with 0.3 M glycine, 0.2% Triton X-100, and 5% normal donkey serum (NDS) for 60 minutes at RT.

For **EdU** detection, cells were processed prior to the blocking step following the manufacturer’s instructions provided in the Click-iT EdU Cell Proliferation Kit (Invitrogen). Primary antibodies were diluted in blocking solution and cells were incubated for overnight at 4°C. Antibodies used in experiments are listed in **Table 1**. After primary antibody incubation, cells were washed three times with TBST (PBS with 0.1% Tween-20) for 5 minutes each and incubated with fluorescently labeled secondary antibodies diluted in TBST solution for 2h at RT. Cells were then washed three times with TBST, counterstained with DAPI, and mounted using ProLong Gold Antifade Mountant (Invitrogen). Images were acquired at 20x magnification using a Keyence BZ-X800 microscope.

**Table 1.** Antibodies used in the experiments.

| REAGENT or RESOURCE | SOURCE | CATALOGUE # |
| --- | --- | --- |
| **Antibodies** |  |  |
| Rabbit anti-DCX<br>(IHC 1:200; WB 1:1000) | Abcam | ab18723 |
| Mouse anti-beta-3 Tubulin<br>(IHC 1:250; WB 1:1000) | Thermo Fischer Scientific | MA1-118 |
| Rabbit anti-Sox2<br>(IHC 1:200; WB 1:1000) | Abcam | ab97959 |
| Donkey anti-rabbit 488<br>(ICC/IHC 1:500) | Jackson ImmunoResearch Labs | 711-545-152 |
| Donkey anti-mouse 488<br>(ICC/IHC 1:500) | Jackson ImmunoResearch Labs | 715-545-151 |
| Donkey anti-rabbit Cy3<br>(ICC/IHC 1:500) | Jackson ImmunoResearch Labs | 711-165-152 |
| Donkey anti-rabbit Cy5<br>(ICC/IHC 1:500) | Jackson ImmunoResearch Labs | 711-175-152 |
| Donkey anti-mouse Cy5<br>(ICC/IHC 1:500) | Jackson ImmunoResearch Labs | 715-175-151 |
| Goat anti-mouse IgG-HRP<br>(WB 1:10000) | Promega | W402B |
| Goat anti-rabbit IgG-HRP<br>(WB 1:10000) | Promega | W4011 |
| Mouse anti-NFH<br>(WB 1:5000) | Proteintech | 60331-1-Ig |
| Mouse GAPDH<br>(WB 1:2500) | Thermo Fischer Scientific | MA5-15738 |
| Rabbit anti-APP<br>(WB 1:1000) | Millipore Sigma | A8717 |
| Rabbit anti-BACE1<br>(WB 1:1000) | Cell signaling | 5606S |
| Mouse anti-AT8<br>(WB 1:1000) | Thermo Fischer Scientific | MN1020 |
| Mouse anti-Tau5<br>(WB 1:1000) | Thermo Fischer Scientific | AHB0042 |
| Rabbit anti-DYRK1A<br>(WB 1:1000) | Cell signaling | 2771S |
| Rabbit anti-BACE2<br>(WB 1:1000) | Cell signaling | 80357 |
| Mouse anti-SOD1<br>(WB 1:1000) | Santa Cruz | sc-17767 |
| Rabbit anti-S100β<br>(WB 1:1000) | Abcam | ab52642 |

### Western Blotting

Cells were washed three times with ice-cold 1X PBS and lysed on ice for 10 minutes using RIPA buffer supplemented with protease and phosphatase inhibitor cocktails (Thermo Fisher Scientific). Lysates were sonicated on ice at 20% power for three cycles of 15 seconds each, with 5-second rests between sonication. Following sonication, samples were centrifuged at 10,000 x g for 15 minutes at 4°C to pellet insoluble debris, and the clarified supernatant was collected. Protein concentrations were determined using the Pierce BCA Protein Assay Kit (Thermo Fisher Scientific). For electrophoresis, equal amounts of protein were mixed with NuPAGE LDS Sample Buffer and NuPAGE Reducing Agent (Invitrogen), then denatured by boiling at 95°C for 5 minutes. Samples were resolved on Bolt Bis-Tris Plus SDS-PAGE gels (Thermo Fisher Scientific) using Bolt MES SDS Running Buffer. Proteins were transferred to nitrocellulose membranes using the iBlot 3 Dry Blotting System (Invitrogen).

Membranes were blocked for 1 hour at room temperature in 5% non-fat dry milk prepared in Tris-buffered saline with 0.1% Tween-20 (TBST). Primary antibodies diluted in 5% milk/TBST were incubated with membranes overnight at 4°C. Antibodies used in experiments are listed in **Table 1**. After washing membranes three times for 15 minutes each in TBST, membranes were incubated with horseradish peroxidase (HRP)-conjugated secondary antibodies diluted in 5% milk/TBST for 2 hours at room temperature. Following secondary antibody incubation, membranes were washed three times as before and developed using SuperSignal West Pico PLUS Chemiluminescent Substrate (Thermo Fisher Scientific). Blots were imaged with the Azure Biosystems 300q Image Western Blot Imaging System.

Densitometric quantification of band intensities was performed using Fiji (NIH). Protein expression levels were normalized to housekeeping controls such as GAPDH.

### Enzyme-Linked Immunosorbent Assay (ELISA) for amyloid-beta in neuronal conditioned media

Conditioned media (CM) were collected from 10 weeks old neuronal cultures derived from iPSCs. For CM samples, cells were rinsed three times with Dulbecco’s phosphate-buffered saline (DPBS) to remove residual media, then cultured in 1 mL of fresh media for 48 hours. Following incubation, conditioned media were harvested and snap-frozen in liquid nitrogen and stored until analysis.

To quantify amyloid-beta isoforms, Human β Amyloid (1-40) ELISA Kit Wako II (Fujifilm Irving, 298-64601) and Human β Amyloid (1-42) ELISA Kit Wako (Fujifilm Irving, 298-62401) were used according to the manufacturer’s protocols. Conditioned media samples were diluted 1:4 for Aβ40 measurement and 1:2 for Aβ42 measurement before loading. All ELISA plates were read on an microplate reader at the 450nm wavelength, and data were processed following kit guidelines. Resulting amyloid-beta concentrations were normalized to the total protein extracted from the corresponding wells.

## STATISTICS

In all graphs, data is shown as mean ± SEM. Prism (Graphpad) was used for statistical analysis with tests indicated in figure legends. The following was used for p values: *p < 0.05, **p < 0.01, ***p < 0.001, and ****p < 0.0001.

## ACKNOWLEDGEMENTS

Graphical abstract, Figure 1A, 1B, and 5A were created Biorender.com. This work was financially supported by National Institutes of Health (NIH), National Institute on Aging (NIA) AG079002, AG057468, AG076940, and AG033570 (OL).

